# Strong positive allometry of bite force in leaf-cutter ants increases the range of cuttable plant tissues

**DOI:** 10.1101/2022.09.28.509980

**Authors:** Frederik Püffel, Flavio Roces, David Labonte

**Affiliations:** Department of Bioengineering, Imperial College London, UK; Department of Behavioural Physiology and Sociobiology, University of Würzburg, Würzburg, Germany

## Abstract

*Atta* leaf-cutter ants are the prime herbivore in the Neotropics: differently-sized foragers harvest plant material to grow a fungus as crop. Efficient foraging involves complex interactions between worker-size, task-preferences and plant-fungus-suitability; it is, however, ultimately constrained by the ability of differently-sized workers to generate forces large enough to cut vegetation. In order to quantify this ability, we measured bite forces of *A. vollenweideri* leaf-cutter ants spanning more than one order of magnitude in body mass. Maximum bite force scaled almost in direct proportion to mass; the largest workers generated peak bite forces 2.5 times higher than expected from isometry. This remarkable positive allometry can be explained via a biomechanical model that links bite forces with substantial size-specific changes in the morphology of the musculoskeletal bite apparatus. In addition to these morphological changes, we show that bite forces of smaller ants peak at larger mandibular opening angles, suggesting a size-dependent physiological adaptation, likely reflecting the need to cut leaves with a thickness that corresponds to a larger fraction of the maximum possible gape. Via direct comparison of maximum bite forces with leaf-mechanical properties, we demonstrate (i) that bite forces in leaf-cutter ants need to be exceptionally large compared to body mass to enable them to cut leaves; and (ii), that the positive allometry enables colonies to forage on a wider range of plant species without the need for extreme investment into even larger workers. Our results thus provide strong quantitative arguments for the adaptive value of a positively allometric bite force.

## Introduction

Leaf-cutter ants are remarkable in many ways: They are the world’s first farmers; their colonies collect plant material to grow a fungus as crop – a mutualism that originated around 50 million years ago [1, 2]. They are considered the principal herbivore in the Neotropics; *Atta* ants are estimated to consume about 15 % of the foliar biomass produced by Neotropical trees [3, 4]. And last, they display a level of polymorphism, linked to size-dependent task-preferences and a complex ecology, exceptional even within social insects [e. g. 5, 6].

Among the diverse tasks arising in a leaf-cutter ant colony are brood care, fungus gardening, worker transport, soil excavation, nest defence and foraging [2, 5, 7]. In particular foraging behaviour has received considerable attention, as it is central to leaf-cutter ant ecology [e. g. 5, 8–34]. Key to foraging success is the cutting of small fragments from vegetation – the energetically most demanding task faced by the colony [35]. Although many aspects of foraging involve complex biological interactions [e. g. 36], cutting plant fragments is, to first order, a mechanical problem: The ability to cut a leaf or fruit is determined by the maximum available bite force, the morphology of the mandible, and the material and structural properties of the plant. Perhaps surprisingly, this mechanical foundation of leafcutting has received comparatively little quantitative attention [e. g. 37–39]. Larger workers possess larger mandible closer muscles and may thus be reasonably expected to produce larger bite forces, which may in turn enable them to cut a wider range of plant materials. Indeed, they tend to be more likely to cut tougher and denser leaves [5, 8, 40–43]. Understanding the exact relation between bite force and worker size is crucial for the analysis of leaf-cutter ant foraging behaviour, because it holds the key to distinguish between foraging assignments based on ability (which ants can cut a given leaf) vs. suitability (which ants should cut a given leaf).

This distinction is relevant for assessing the evolution and possible adaptive advantage of worker polymorphism, and for predictions on ‘optimal’ foraging strategies based on ergonomic considerations [e. g. 8, 30, 43].

We previously predicted the scaling of bite force based on the morphology of the bite apparatus in *Atta vollenweideri* leafcutter ants [44]. However, direct experimental confirmation of this prediction is missing and difficult to obtain, because ants are small. To make matters worse, bite forces in insects generally depend on the mandibular opening angle [45–47], which is challenging to control experimentally. Here, we address both difficulties with the help of a custom-built bite force set-up and first principles biomechanical analysis. We report direct bite force measurements for ant workers spanning more than one order of magnitude in body mass, relate our results to the foraging behaviour of leaf-cutter ant colonies, and discuss both the magnitude and scaling of bite forces in a comparative framework.

## Materials & methods

### Study animals

Bite force experiments were conducted with *A. vollenweideri* ants from three colonies, all founded and collected in Uruguay in 2014 (see Fig. 1A). The colonies were kept at 25°C and 50 % humidity in a climate chamber under a light:dark cycle of 12 h:12 h, and were fed with fresh bramble leaves, cornflakes and honey water *ad libitum*. We collected around 80 workers from the foraging areas of each colony (n = 248), covering approximately the entire size range (1 - 50 mg). We excluded minims (body mass < 1 mg), because their bite apparatus is morphologically distinct [N Imirzian, F Püffel and D Labonte, in preparation 48], and they do not generally partake in foraging [5].

**Figure 1.**
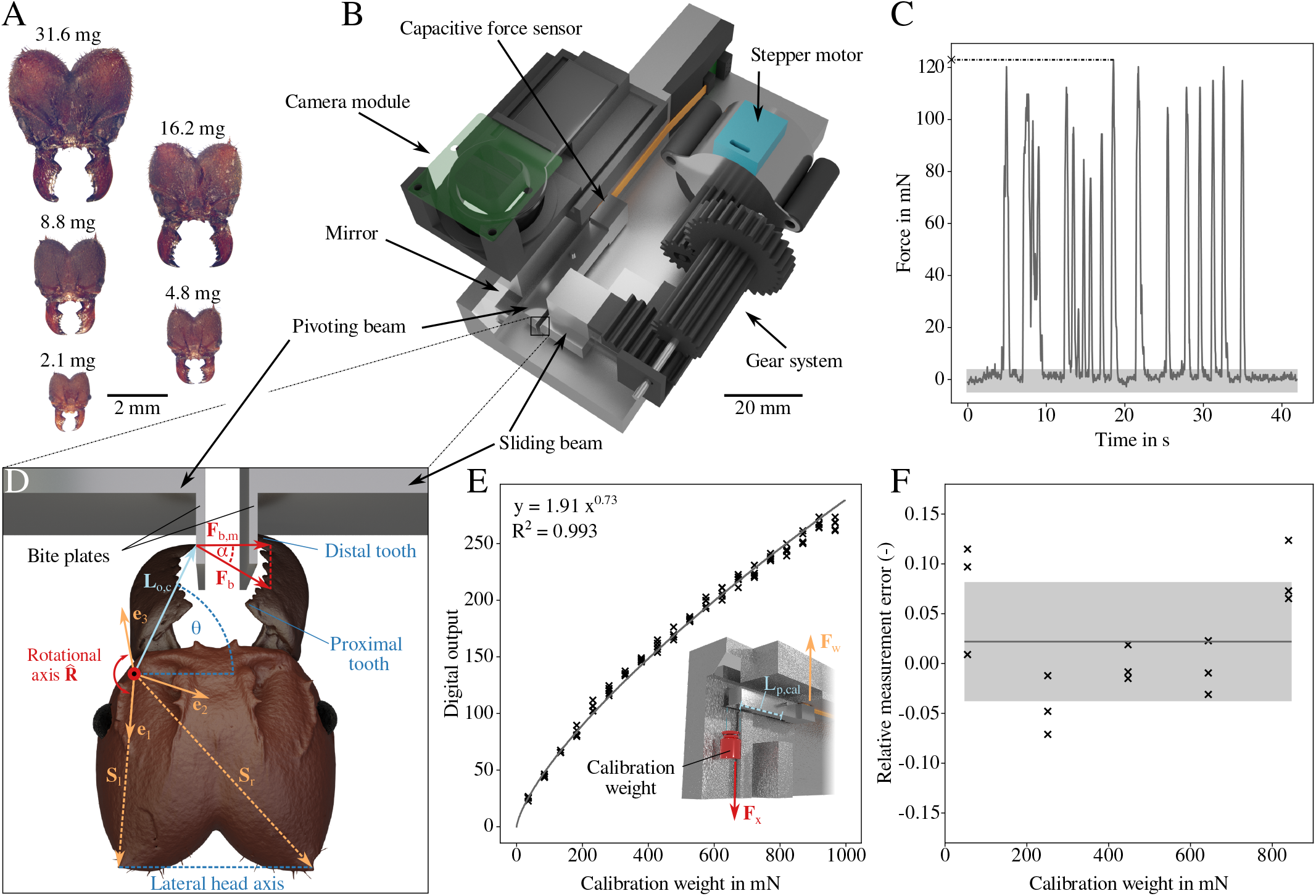
(**A**) We measured bite forces of polymorphic *Atta vollenweideri* leaf-cutter ants ranging between 1.5 and 46.8 mg in body mass. (**B**) To measure bite forces, we built a custom-designed force setup based on a capacitive force sensor and a lever mechanism. The ants bit onto two bite plates, one protruding from a pivoting beam connected to the force sensor, the other protruding from a sliding beam. The distance between the two bite plates can be varied by moving the sliding beam via a geared stepper motor. Bite experiments were filmed with a top-down camera, which also recorded a side view from a 45° mirror. (**C**) When placed in front of the bite plates, ants readily bit. The measured forces exceeded the sensor noise (shaded area) by at least a factor of eight. From each force trace, the maximum force (cross) was extracted for further analysis. (**D**) From the video recordings, we further extracted the coordinates of the mandible joint centre, head spikes **S**_*l*_ and **S**_*r*_, outlever length **L**_*o,c*_, and the most distal and proximal tooth tips (for more details, see SI). The rotational axis of the mandible 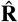 was projected onto the local head coordinate system (**e**_1_, **e**_2_, **e**_3_). The mandibular opening angle *θ* was defined as the angle between the lateral head axis, span by the head spikes, and the projection of the largest outlever onto the plane of rotation. The magnitude of the bite force |**F**_*b*_| was extracted from the measured force |**F**_*b,m*_| and the misalignment angle *α*, extracted from the video recordings, as defined by Eq. 1. For simplicity, the depicted ant bites with its largest outlever, i. e. with the tip of the most distal tooth **L**_*o,c*_ = **L**_*o,d*_, and all vectors are shown in the plane of rotation, i. e. they indicate the effective outlever, **L**_*o,e f f, c*_ (for more details, see SI). The head orientation during experiments however may be different such that the angles *θ* and *α* were not measured directly from the recordings, but were inferred indirectly via their vector-algebraic basis using 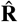 and the local coordinate system (**e**_1_, **e**_2_, **e**_3_; see text). (**E**) For sensor calibration, we suspended weights between 5 and 100 g from the bite plate of the pivoting lever at a distance *L*_*p,cal*_ to the pivot. A linear regression on log_10_-transformed data characterised the relationship between measured output and force with high accuracy (R^2^ = 0.993). (**F**) A subsequent error analysis using intermediate weights between 7 - 87 g yielded relative errors between measured and expected forces of 2 % (accuracy, black line) with a standard deviation of 6 % (precision, shaded area), independent of weight (see text).

### Experimental set-up

Bite forces were measured with a custom-built set-up based on a capacitive force sensor (SingleTact S8-1N, Pressure Profile Systems, Inc., California, USA, data acquisition frequency 33 Hz, maximum force 1 N, resolution 2 mN). In order to distribute the force equally across the sensor area, we used a lever mechanism to convert *point*-like bite forces into an *areal* compression (see Fig. 1B). Ants bit onto two thin bite plates (1 mm long, and ≈ 0.15 mm thick, see Fig. 1D), protruding from two mechanically uncoupled beams (6 × 6 × 40 mm and 6 × 12 × 6 mm, respectively). Both beams and the terminating bite plates were manufactured in one single piece from stainless steel to minimise compliance, using electrical discharge machining. The beam that transmits the bite force to the sensor was connected to a metal base plate, made from a single aluminium block using CNC milling, via a small hinge placed in the beam centre. As an ant bites onto the bite plates, the opposite end of this pivoting beam is pressed onto the sensor, which is glued to a vertical wall protruding from the base. At the maximum load of 1 N, the sensor only deforms approximately 10 *μ*m [manufacturer data]. Because the gear ratio of the pivoting beam is unity, the magnitude of this deformation is equivalent to the maximum displacement of the bite plate arising from beam rotation. As bite plate bending is also negligible (see SI), we measured approximately isometric bite forces. The second beam, in turn, was connected to a lubricated rail, so that its position relative to the pivoting beam can be altered via a stepper motor (28BYJ-48, 5 V), connected to a gearing system 3D printed from PLA. This design allowed us to vary the distance between the two bite plates by less than 100 *μ*m per motor step. The minimum required gape to bite both bite plates, equivalent to the shortest distance between their outer surfaces, was 0.5 mm, and the largest gape used during experiments was 3.0 mm. This range is similar to the range of head widths of the ants used in this experiment (1.1 - 4.5 mm).

In order to extract coordinates of key landmarks (see below), a top-down camera was synchronised to the sensor recording, and recorded the bite experiments at 30 fps, (camera module v 2.1, 8 MP, Raspberry Pi Foundation, Cambridge, UK; with a Black Eye HD macro lens, Eye Caramba Ltd., Helsinki, Finland). A mirror, tilted 45° with respect to the camera plane, provided depth information. Force sensor, motor, camera and a touchscreen (7 inch HDMI-LCD Display, Elecrow, Shenzhen, China) were connected to a Raspberry Pi (v 3B+, Raspberry Pi Foundation, Cambridge, UK). The system was operated via a custom software written in Python [v 3.5, see 49, and SI for details on the user interface and circuit].

### Sensor calibration, accuracy and precision

The sensor was calibrated using a series of calibration weights (Kern & Sohn GmbH, Balingen, Germany), suspended from the pivoting lever via a thin cotton thread (the weight of the cotton thread was small, ≈ 0.015 g, and hence neglected). The thread was placed at around one-quarter of the length of the bite plate measured from its base, resulting in a moment arm around the pivot of *L*_*p,cal*_ ≈ 20.25 mm. The set-up was rotated such that the measurable force vector aligned with gravity (see Fig. 1E). We used 20 weights between 5 and 100 g; a weight of 1.4 g was required to establish equilibrium around the pivot, and was consequently subtracted from all readings. The range of effective weight forces was hence 35 - 970 mN, approximately spanning the range of measured bite forces. Each weight was suspended five times in random order, and the steady-state sensor reading averaged across ≈ 2 s was used for sensor calibration. The relationship between weight and sensor output was characterized via an Ordinary Least Squares regression on log_10_-transformed data in order to prioritise minimisation of relative error over absolute error; the high R-squared of the regression line indicates robust calibration (R^2^ = 0.993, see Fig. 1E).

To quantify calibration accuracy and precision, five additional weights (7 - 87 g) were measured three times each, and the relative error was calculated as *ε* = 1 *−* |**F**_*w*_|*/*|**F**_*x*_|, where **F**_*x*_ is the predicted and **F**_*w*_ is the measured force, respectively (see Fig. 1F. Throughout this text, we write vectors as bold; unit vectors are labelled with a hat as in 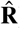). The average measurement error, the sensor accuracy, was 2 ± 6 % independent of weight [Linear Mixed Model (LMM) with weight as fixed and repetition as random effect: 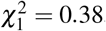, p = 0.53; the average magnitude of the measurement error was 5 ± 4 %]; the standard deviation indicates the precision of the measurements.

To quantify sensor noise, the output of the unloaded sensor was recorded for 100 s. Discrete Fourier transformation suggested a constant power spectral density, indicating white noise [50, p. 61] This noise had a standard deviation of 1 mN and a range of -5 to 4 mN. The maximum positive value of this range was about eight times smaller than the smallest measured bite force (see Fig. 1C). Sensor drift was significant [Spearman’s correlation coefficient: *r*_*s*,2867_ = -0.26, p < 0.001; see manufacture data sheet], but small enough to be inconsequential across the time scale of our measurements (cf. 1 mN/min vs ≈30 - 60 s).

In a last validation step, we directly compared measurements with our set-up with those of an established set-up based on a substantially more expensive piezoelectric sensor [51, 52]. To this end, maximum bite forces of three house crickets (*Acheta domesticus*) were measured with three repetitions on both setups, by two independent operators each. A LMM with set-up and operator as fixed and specimen as random effect showed no significant influence of neither set-up nor operator on bite forces [LMM versus random intercept model: 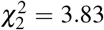, p = 0.15; set-up: t_31_ = 1.68, p = 0.10; operator: t_31_ = 0.95, p = 0.35].

The camera was calibrated assuming an inverse size-distance relationship, *L*_2_*S*_2_ = *L*_1_*S*_1_, where *S*_1_ and *S*_2_ are the apparent sizes of an object at distances *L*_1_ and *L*_2_ to the camera aperture. The apparent size of a calibration target (5 units on 1 mm-grid paper), placed on the top surface of the metal base at a distance *L*_1_ ≈ 32 mm to the camera, was measured from ten images, and the pixel-to-mm conversion factor *C*_1_ was extracted. This conversion factor changes approximately in direct proportion to the camera distance as *C*_2_ = *C*_1_*L*_2_*/L*_1_, where *L*_2_ is the distance between the camera and the imaged object. In contrast to *L*_1_, *L*_2_ is not constant, but varies with the vertical position of the object coordinate, which was extracted using the mirrored side view.

For validation, the grid paper was photographed at 25 different positions along the bite plate, extracted from five trials, and its physical size was calculated from the calibration. The relative error was 1±2 %. This error changed significantly with grid paper-lens distance – the physical dimensions were increasingly underestimated at smaller distances [LMM with lens distance as fixed effect and trial as random effect: 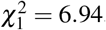, p < 0.01]. This effect, however, was miniscule (1.5 % across the entire bite plate length, 6 mm), corresponding to only ≈ 60 *μ*m, or about 10 % of the smallest measured mandible length; it is thus con-sidered negligible.

### Experimental protocol

To measure bite forces, individual ants were held in front of the bite plates using insect tweezers. Ants were eager to bite, typically executing numerous bite cycles in quick succession (see Fig. 1C). After completing at least five bite cycles, or exceeding a total bite duration of 10 s, the measurement was terminated, each ant was weighed (AX304 Microbalance, 310 g x 0.1 mg, Mettler Toledo, Greifensee, Switzerland) and isolated, and the maximum force was extracted. To obtain bite forces from a maximum range of sizes and varying opening angles without pseudoreplication, we measured each ant only once at a single bite plate distance. In practice, we gradually increased the bite plate distance from 0.5 to 3.0 mm; for each distance, we used ants of all sizes capable of biting onto the plates. For small plate distances, this included almost all ants; for the largest plate distance, however, only larger specimen were able to bite. We assume that ants bit with maximum muscle activation at all opening angles, as indicated by measurements involving direct muscle stimulation in closely related *Atta cephalotes* [47].

### Extraction of landmarks

In order to obtain the opening angle, the orientation of the bite force vector, and the length of the mandible outlever (see below), we extracted the coordinates of a series of landmarks from the video frame corresponding to the time of maximum bite: (i) the contact point between mandible and bite plate; (ii) the tip of the most distal and proximal mandibular teeth; (iii) the mandible joint centre (defined as in [44]); and (iv) a pair of distinct head spikes (see Fig. 1D and SI figure). In addition, we extracted the orientation of the mandible joint axis 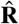 via kine-matics analysis of the mandible motion [47, 53, V Kang, F Püffel and D Labonte, in preparation]. In order to position 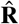 on the biting ant, we introduced a local head coordinate system (**e**_1_, **e**_2_, **e**_3_) based on the vectors connecting the joint centre with the tips of both head spikes, **S**_*l*_ and **S**_*r*_, respectively. The coordinate axes were defined as **e**_1_ = **S**_*l*_*/*|**S**_*l*_|, **e**_3_ = (**S**_*l*_ × **S**_*r*_)*/*|**S**_*l*_ × **S**_*r*_|, and **e**_2_ = **e**_3_ × **e**_1_. The estimated rotational axis was then projected onto this local head coordinate system (for more details, see SI). In line with previous work [54], we define the opening angle as the angle between the lateral head axis and the largest effective outlever. The lateral head axis was defined as the vector connecting both head spikes (**S**_*r*_ *−* **S**_*l*_, see Fig. 1D); the largest effective outlever was defined as the length of the vector which connects the tip of the most distal mandible tooth and the joint rotational axis in the plane of rotation, 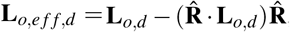, where **L**_*o,d*_ is any vector which connects the most distal tooth tip and the rotational axis (see SI for more details). The opening angle then follows from basic vector algebra as *θ* = arccos[(**L**_*o,e f f,d*_ · (**S**_*r*_ − **S**_*l*_))/(|**L**_*o,e f f,d*_||**S**_*r*_ − **S**_*l*_|)].

### Confounding effects due to size-differences

Measuring bite forces across a large size range poses at least two challenges. First, the signal-to-noise ratio typically decreases with animal size, as smaller animals generally bite with less force. We addressed this challenge by selecting a force sensor capable of measuring small and large bite forces with sufficiently high resolution (see above). Second, the characteristic dimensions of the set-up are relatively larger for small animals. The implications of this size-effect are perhaps less obvious and, unfortunately, more complex. To appreciate the problem, consider a small ant of 3 mg body mass biting onto bite plates that are 1 mm apart. This bite plate distance is approximately equal to the distance between both mandible joints; a bite with the distal-most teeth would thus involve a mandibular opening angle of about 90°. For a bite with the most proximal teeth, in turn, the opening angle is larger, around 110° (see Fig. 1A), and the effective outlever is smaller. Consider next a large ant of 45 mg biting onto the same bite plates. For a distal bite, the bite plate distance corresponds to only 40 % of the joint distance, and thus a much lower opening angle of about 70°. Notably, the ant’s ability to bite with proximal parts of her mandible is limited because the length of the mandible blade exceeds the length of the bite plate by about 20 % (see Fig. 1D). Evidently, for a given configuration of the experimental setup, both the range of possible mandibular opening angles and bite positions along the mandible blade vary systematically with size. Because both directly affect the magnitude of bite force [see Eq. 3 and 47], an unbiased comparison of bite force magnitude across animal sizes, requires four corrective steps, implemented here with the ultimate aim to extract the maximum bite force an ant worker can produce when biting with an equivalent point of her mandible (the tip of the most distal mandible tooth. See Fig. 2). First, we correct for the misalignment between *measurable* force vector, **F**_*b,m*_, and *applied* bite force vector, **F**_*b*_. Second, we correct for differences in moment arms around the bite lever pivot, arising from variation of the mandible contact point along the bite plate. Third, we account for variation in bite position along the mandible blade, which alters the mechanical advantage of the force transmission system. Fourth, we account for differences in mandibular opening angle.

**Figure 2.**
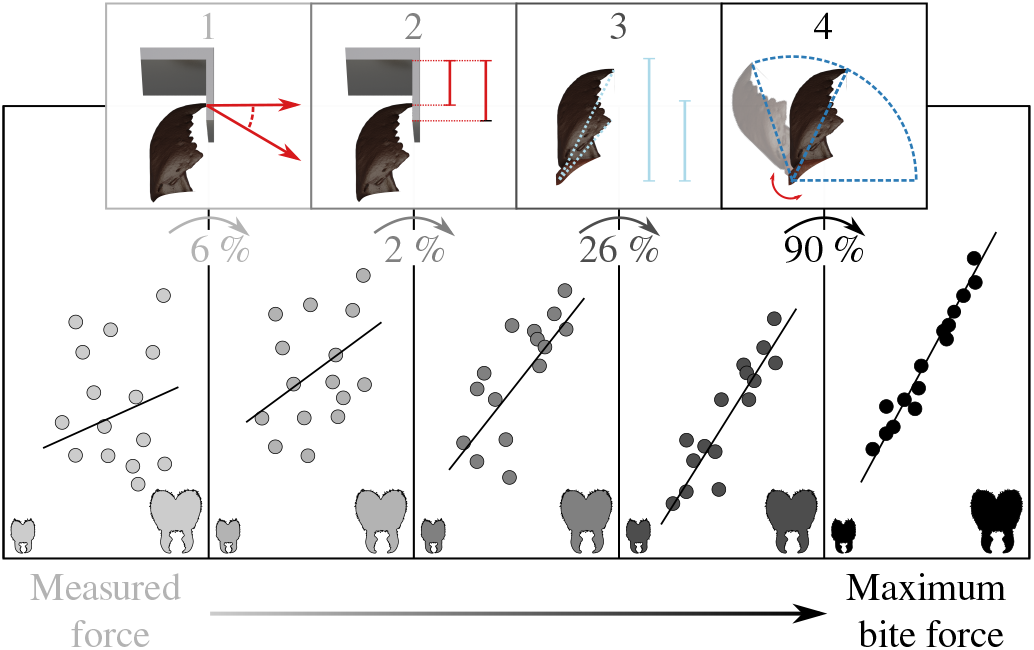
In order to extract the maximum bite force from the measured data, we account for four confounding effects: First, we account for the misalignment between bite force orientation and the force-sensitive axis. Second, we account for variation in bite contact point on the bite plate, which results in different moment arms. Third, we account for differences in bite position along the mandible blade, which determines the mechanical advantage of the musculoskeletal force transmission system. Fourth, we account for variation in mandibular opening angle, which is associated with variation in mechanical advantage, fibre length and pennation angle, all of which influence bite forces. Implementation of these corrections reduces the size-dependent variation introduced by the experimental design and animal behaviour, and thus enables an unbiased comparison of maximum bite force capacity across animals of different sizes. The data depicted in this schematic is not ‘real’ data, but merely serves for illustration. The ‘actual’ relative change in force caused by each correction, averaged for all ants, is shown in percent.

The first correction is necessary, because the sensor measures 1D compression, **F**_*b,m*_, but the applied bite force vector, **F**_*b*_, may deviate from this line of action by a misalignment angle *α* (see Fig. 1D). The orientation of **F**_*b,m*_ is approximately equal to the plane normal of the bite plate; the orientation of the bite force vector is defined by the cross product between mandible outlever **L**_*o,c*_ and the rotational axis, where **L**_*o,c*_ is any vector connecting the joint axis of rotation and the bite contact point [for more details, see 47]. The resulting correction factor reads 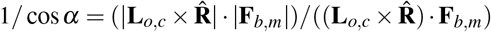 [for more details, see 47]. This correction was typically larger for larger animals, which more frequently bit at opening angles below 90° (see results), leading to larger misalignment angles.

The second correction is necessary, because the point of force application on the bite plate itself may vary across ants – smaller ants have shorter mandibles and are thus more likely to bite at the end part of the bite plates (see Fig. 1D). For each bite, we thus measured the lever arm around the bite lever pivot, *L*_*p*_, defined as the distance between the bite contact point and the pivot, measured in the plane of rotation. To remove variation due to *L*_*p*_, we introduce the correction term Γ = *L*_*p,cal*_*/L*_*p*_, where *L*_*p,cal*_ is the distance between the pivot and the point of application of weights during sensor calibration. Γ was always close to 1, as the length of the bite plate is small in comparison to *L*_*p,cal*_ (≈ 5 %); the effect of this correction was thus miniscule.

The third correction is necessary, because the bite force may be transferred onto the bite plate at an arbitrary position along the mandible; an ant biting with the distal end of its mandible may use the same muscle effort but nevertheless produce a smaller measured bite force than an ant biting with the proximal end of its mandible, because the mechanical advantage of the force transmission system differs [see e. g. 55]. To correct for this variation, we converted the measured bite force into an equivalent bite force at a fixed point on the mandible blade (the most distal tooth tip). To this end, we extracted the length of the vector which connects the contact point between mandible and bite plate with the joint rotational axis in the plane of rotation, **L**_*o,e f f,c*_ – the effective outlever of the bite force measurement. We then calculated the bite force for an hypothetical bite exerted via the tip of the most distal tooth, characterised by an effective outlever **L**_*o,e f f,d*_ [see above and 47, for more details].

After implementation of corrections (i-iii), the magnitude of the bite force |**F**_*b,θ*_ | follows as:

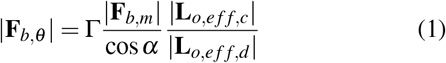

A fourth and last correction is then necessary, because bite forces vary systematically with mandibular opening angle [46, 47, 56–59]. In general, the bite force at any opening angle may be written as the product between muscle stress *σ*, the physiological cross-sectional area of the mandible closer muscle *A*_*phys*_, the average pennation angle *φ* of the muscle fibres, and the mechanical advantage of the mandible lever system, defined as the ratio between effective inand outlever |**L**_*i,e f f*_ |*/*|**L**_*o,e f f*_ | [see 44, 47, for details]:

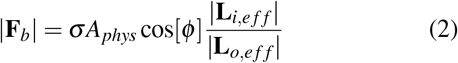

Here, the effective inlever is the length of the shortest vector which connects the line of action of the net muscle force vector with the axis of rotation, and the effective outlever is the length of the vector which connects the rotational axis with the bite contact point on the mandible blade, all defined in the plane of rotation. Equation 2 is in widespread use to model the biomechanics of musculoskeletal lever systems, but a major complexity is only implicit: The muscle stress, the pennation angle, and the mechanical advantage all depend on the mandibular opening angle [46, 47, 58, 60, 61]. The resulting variation in bite force with opening angle can be substantial: in *A. cephalotes* majors, bite forces at small opening angles are around five times larger than those at large opening angles [see 47, and Fig. 2]. In order to estimate maximum bite forces from bite forces measured at different opening angles, it is thus necessary to determine how muscle stress, pennation angle, and mechanical advantage vary with opening angle:

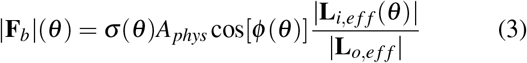

The muscle stress depends on opening angle, because contractions at different opening angles involve different fibre lengths, and muscle has a characteristic force-length dependence [62–64]. The variation of stress with fibre length can be characterised by an empirical function 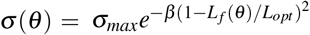, where *β* is a shape parameter, and *L*_*opt*_ is the fibre length at which muscle stress is maximum [64]. We recently demonstrated that the functions cos[*φ* (*θ*)], **L**_*i,e f f*_ (*θ*) and *L*_*f*_ (*θ*) can be accurately predicted from first principles, using morphological reference measurements for the muscle volume, average muscle fibre length, pennation angle, and the effective mandible levers from a single opening angle [see 47, for a detailed derivation and SI for exact mathematical expressions]. In order to obtain these reference measurements, we analysed 13 tomographic scans of *A. vollenweideri* leaf-cutter ants across the size range [for details, see 44].

We assume that all fibres of the closer muscle attach to the tendon-like apodeme via thin filaments [which holds for 98 % of fibres in *A. vollenweideri*, 44], and define the physiological cross-sectional area as the muscle volume divided by the fibre length at which the muscle stress is maximum, *L*_*opt*_ [47]. The maximum muscle stress *σ*_*max*_ = 1.16 MPa and *β* = 5.34 were taken from Püffel et al. [47], where they were determined experimentally for closely related *A. cephalotes* majors. We make the simplifying assumption that these parameters are size-independent, and this assumption is supported by the resulting agreement between theoretical prediction and measured bite force (see below).

The mandibular opening angle *θ*_*opt*_ at which the muscle stress is maximum, *L*_*f*_ (*θ*_*opt*_) = *L*_*opt*_, was then estimated via a nonlinear least squares numerical fitting routine of Eq. 3 in python [v 3.9 49]. *θ*_*opt*_ may be reasonably expected to vary with body size, as it is sensitive to head capsule geometry which changes with size in *A. vollenweideri* [see 44, 48, and N Imirzian, F Püffel and D Labonte, in preparation]. In order to investigate the size-dependence of *θ*_*opt*_, we only fitted bite force measurements from individuals with a body mass that differed by no more than ≈ 25 % from the individuals for which we obtained morphological data from *μ*CT scans. Each size-class bin contained at least eight bite force measurements, except for the largest and smallest ant, which contained two and three bites, respectively.

In order to implement the opening angle correction across worker sizes, we first characterised the size-dependent relationship between bite force and opening angle as outlined above. Next, we calculated a database containing the relationship between predicted bite force, normalised with its maximum, |**F**^*∗*^|, opening angle and size. We rescaled the measured forces, corrected via Eq. 1, |**F**_*b,θ*_ |, to the force expected at an *equivalent* mandibular opening angle – the angle *θ*_*max*_ at which bite forces are maximal 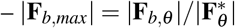, where 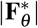 represents the fraction of the maximum bite force produced at *θ*_*max*_. The magnitude of this correction thus depends on both the body mass and the mandibular opening angle of the biting ant; the corrected value used for the final analysis is the maximum bite force the animal can transmit at the distal mandibular tooth (see Fig. 1D).

### Data curation and statistical analysis

We pooled bite force data from all three colonies, because the relationship between bite force and body mass was independent of colony both before and after correcting for differences in opening angle [Analysis of covariance (ANCOVA) on log_10_-transformed data, before correction: F_2,242_ = 1.26, p = 0.28; after correction: F_2,242_ = 0.85, p = 0.43]. Out of 248 bite force measurements, three slightly exceeded the range of calibration forces; two were lower and one higher, but all within 15 % to the nearest calibration force. These measurements neither appeared to be outliers, nor altered the scaling relationship (see results), and were hence kept for the analysis. Scaling relationships were characterised with Ordinary Least Squares (OLS) and Reduced Major Axis (RMA) regression models on log_10_-transformed data in R [v 4.1, 65]. For simplicity, we only report the results of the OLS regressions in the text; the RMA regression results are provided in the SI. Scaling coefficients – the slopes of the regression lines – tend to be higher for RMA regressions [also see 66]; however, the main conclusions of this study are supported by both.

## Results

### Leaf-cutter ant bite forces increase with strong positive allometry

We measured bite forces of *A. vollenweideri* leaf-cutter ants spanning more than one order of magnitude in body mass, m. Measured bite forces |**F**_*b,m*_| ranged from a minimum of 31 mN to a maximum of 1029 mN, and were proportional to m^0.79^ [OLS 95 % CI: (0.74 | 0.85), R^2^ = 0.75, see SI figure], suggesting positive allometry (i.e. a scaling relationship steeper than predicted by isometry, m^0.67^). After correcting for force orientation, and bite position along the bite plate and mandible blade, respectively, bite forces, |**F**_*b,θ*_ |, ranged between 19 and 901 mN (see Fig. 2 and Eq. 1), and were proportional to m^0.85^ [OLS 95 % CI: (0.79 | 0.92), R^2^ = 0.74], in substantial excess of the scaling coefficient of |**F**_*b,m*_| (see Fig. 3B). However, this scaling relationship is still influenced by systematic differences in mandibular opening angle, which, as expected, were systematically smaller for larger animals [Analysis of variance (ANOVA) on log_10_-transformed data: F_1,246_ = 15.24, p < 0.001]. In order to correct for the systematic differences in opening angle, we first assessed the size-dependence of the opening angle, *θ*_*opt*_, at which muscle stress is maximum [see methods and 47]. Notably, *θ*_*opt*_ decreased significantly with body mass [OLS: slope = -0.09, 95 % CI: (−0.15 | -0.04), p < 0.01, R^2^ = 0.56]: *θ*_*opt*_ ≈ 70° for a 1.5 mg ant, but *θ*_*opt*_ ≈ 50° for a 45 mg ant. Next, we calculated the opening angle at which the bite force is maximum, *θ*_*max*_. *θ*_*max*_ may differ from *θ*_*opt*_, because it is the opening angle at which the product between the mechanical advantage, the pennation angle and the muscle stress, as opposed to just muscle stress, is maximal [see eq. 3, 47]. *θ*_*max*_ decreased significantly with size [OLS: slope = -0.06, 95 % CI: (−0.10 | -0.03), p < 0.01, R^2^ = 0.59, represented by the crosses in Fig. 3A]; the bite forces of a 1.5 mg and 45 mg ant are maximal at around 65° and 50°, respectively. Thus, the difference between *θ*_*max*_ and *θ*_*opt*_ is small, supporting our earlier conclusion that the musculoskeletal bite apparatus of leaf-cutter ants has a morphology which maximises the magnitude of the peak bite force [see 47].

**Figure 3.**
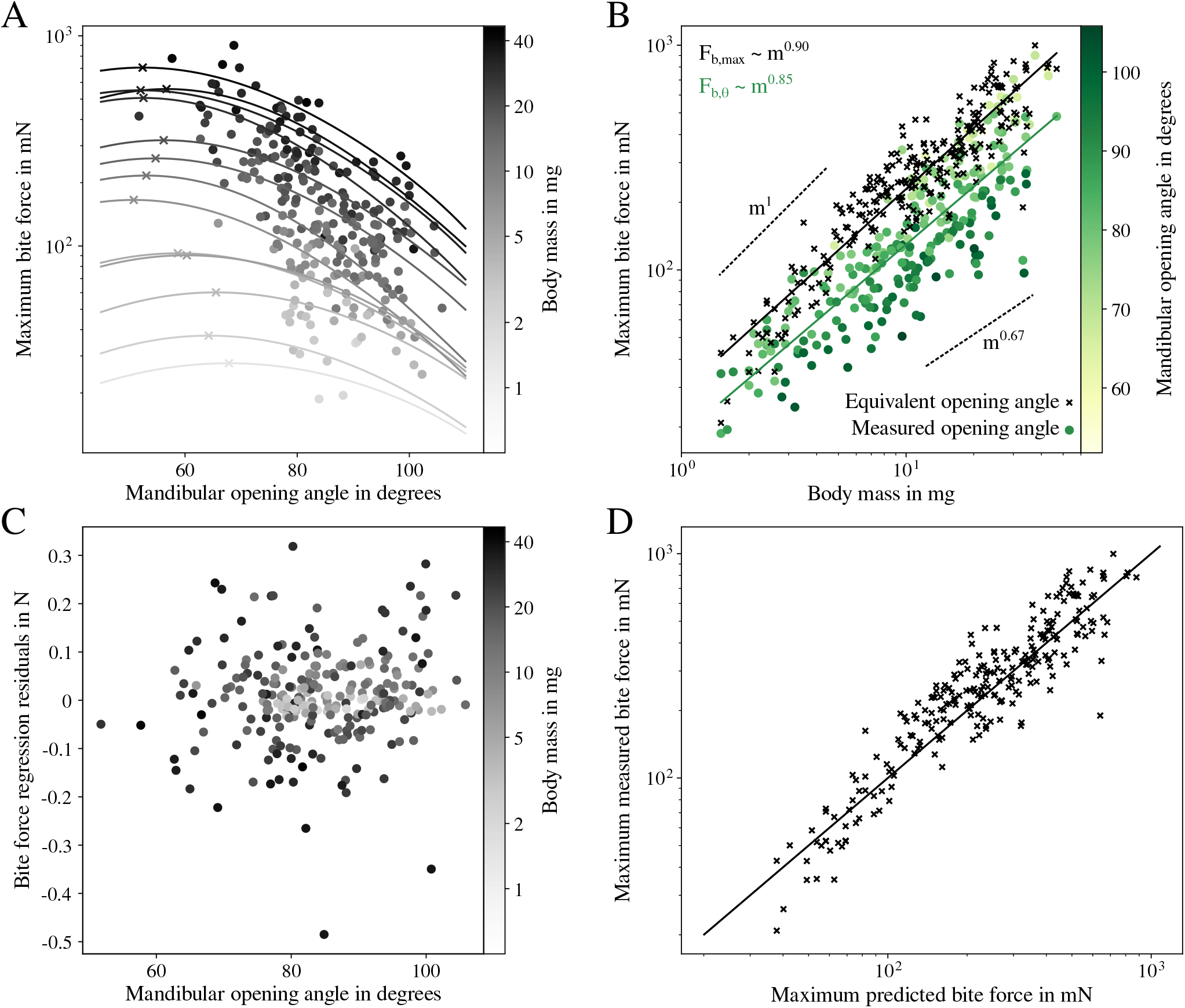
(**A**) Bite forces of *Atta vollenweideri* workers vary with both mandibular opening angle and body mass. Because ants bite onto two bite plates of finite thickness during bite force measurements (see Fig. 1B & D), bites from smaller ants were measured with larger average opening angles (see also Fig. 2 and methods). In order to correct for this systematic difference prior to assessing the scaling of bite force with body mass, the relation between bite force and opening angle for workers of different body masses was predicted via biomechanical modelling (Eq. 3, solid lines), and the maximum bite force was extracted for subsequent scaling analyses (crosses; for details on the correction, see methods). Note that the maximum bite force occurs at systematically larger opening angles for small workers [Ordinary Least Squares (OLS) regression: slope = -0.09, 95 % CI: (−0.15 | -0.04), p < 0.01]. (**B**) Bite forces grew in almost direct proportion to body mass (upper dashed line), in substantial excess of the prediction from isometry (lower dashed line). The effect of the correction for differences in opening angle is a significant increase of the slope and intercept of the regression line, and of the variation explained by body mass (circles represent uncorrected and crosses opening-angle corrected data). (**C**) The residuals of the regression of the opening-angle corrected bite forces against mass (black line and crosses in **B**) are independent of mandibular opening angle [ANOVA: F_1,246_ = 2.43, p = 0.12], supporting the validity of the biomechanical modelling. (**D**) Experimentally measured and corrected bite forces are in excellent agreement with a theoretical prediction via biomechanical modelling (see Eq. 3), as demonstrated by the identity function (solid line). We note that predicted and corrected forces are not strictly independent; the optimum opening angle, affecting the force prediction, was fitted using the measured bite forces (for further details, see methods).

Equivalent bite forces at *θ*_*max*_ and the most distal bite point scale with an even stronger positive allometry, |**F**_*b,max*_| ∝ *m*^0.90^ [OLS 95 % CI: (0.86 | 0.95), R^2^ = 0.86]. The increase in scaling coefficient is surprising, because larger ants bit at significantly smaller opening angles; the intuitive expectation is thus that the bite forces of small ants were underestimated, and the scaling coefficient overestimated. However, this argument neglects the significant decrease of *θ*_*max*_, which reverts the effect: because *θ*_*max*_ decreases more quickly than the opening angle increases, larger ants were measured at less favourable opening angles, and the scaling coefficient was underestimated.

Together, the four corrections result in a significant increase of the scaling coefficient by 0.11, or about 15 % [LMM with correction as fixed and sample number as random effect: 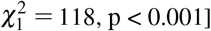, and in an increase of the R^2^ from 0.75 to 0.86. The difference in scaling coefficient may appear small, but this impression is misleading: the difference in maximum bite force between ants varying in body mass by a factor of 30 is 30^0.90^ ≈ 22 compared to 30^0.79^ ≈ 15 – a drop by about 30 %. Clearly, a careful analysis of bite force measurements is required in order to draw meaningful conclusions on scaling relationships.

## Discussion

Leaf-cutter ants are an ecologically and economically important herbivore [4, 6], foraging on a wide variety of plants with different mechanical properties [40, 41]. Key to foraging success is the ability of workers to produce bite forces sufficiently large to cut plant tissues [67]. How does this ability vary with worker size? In order to address this question, we measured the maximum bite forces of *A. vollenweideri* leaf-cutter ants spanning more than one order of magnitude in body mass. In the following discussion, we (i) connect the bite force allometry with its morphological and physiological determinants; (ii) discuss the magnitude and scaling of maximum bite forces in the context of foraging ecology; and (iii) place our findings in an evolutionary and comparative context.

### Bite force allometry can be accurately predicted from muscle architecture and head morphology

Maximum bite forces of *A. vollenweideri* workers show strong positive allometry, |**F**_*b,max*_| ∝ *m*^0.90^. Due to the large variation in worker size, the effect of this positive allometry is rather extreme: the largest ant workers generate maximum bite forces about 2.5 times higher than a theoretical isometric worker with the same body mass, (45*/*1)^0.90*−*0.67^ ≈ 2.5. Can this extreme positive allometry be understood from the morphology and physiology of the *Atta* bite apparatus?

We have previously extracted the relevant morphological bite force determinants in *A. vollenweideri* workers from tomographic scans [see 44]. Using these data in conjunction with Eq. 3 led to the prediction that bite forces should scale as m^0.88^ (95 % CI: 0.81 | 0.95), a positive allometry largely driven by a disproportional increase in physiological cross-sectional area of the mandible closer muscle [44]. This prediction is in rather close agreement with our bite force measurements. The small difference stems from the fact that we previously extracted the morphological bite force determinants from scans, where the mandibles were maximally closed, whereas *θ*_*max*_ varies across sizes (see Fig. 3A). In order to account for this variation in our morphological prediction, and to estimate the intercept of the scaling relationship, we use the stress and force-length shape parameter measured for closely related *A. cephalotes* [47], the change in *θ*_*opt*_ as observed in this work, and a biomechanical model which links morphology, opening angle and bite force to directly predict maximum bite forces [see 47, and Fig. 3A].

The maximum bite forces predicted by this calculation are proportional to m^0.91^ [OLS 95 % CI: (0.82 | 1.01), R^2^ = 0.98], with an intercept of 1.42 [units: mN, mg; OLS 95 % CI: (1.32 | 1.52]. Both estimates are almost identical to the results obtained from direct measurements: m^0.90^ (scaling coefficient), 1.46 (intercept) [units: mN, mg; OLS 95 % CI: (1.41 | 1.51)]. In other words, the ratio between maximum measured and predicted bite forces is close to unity (1.11±0.30), and independent of body mass [ANOVA on log_10_-transformed data: F_1,246_ = 0.23, p = 0.63, see Fig. 3D]. We thus conclude that the scaling of bite forces can be predicted to reasonable accuracy from morphological measurements.

Predicting bite performance from morphology is of considerable interest to evolutionary biologists, palaeontologists and biomechanists alike, as *in-vivo* force measurements are often challenging if not impossible to obtain. Consequently, theoretical models to predict maximum bite force have been developed and deployed for numerous taxa [e. g. 47, 61, 68–70]. In arthropods, the two key obstacles facing such theoretical efforts are the extraordinary variation in reported muscle stresses [e. g. 71, 72], and the uncertainty in estimates of the physiological cross-sectional area, which requires knowledge of the optimal fibre length [47]. The fact that we were able to accurately predict both the magnitude and scaling of bite forces in *A. vollenweideri* using physiological parameters measured in closely related *A. cephalotes* suggests that muscle stress and shape parameter *β* are conserved across sizes and within *Atta*, cautiously indicating that the intraspecific and intrageneric variation of bite forces may be predictable based on morphological data alone.

### Positive allometry of bite force has substantial benefits at colony level

Bite force is a non-pareil performance measure which influences access to food sources and high-quality mating partners [73]. As a result, the variation in morphology and physiology of the bite apparatus across species often reflects species-specific needs [45, 56, 57, 59, 74]. What needs have shaped the evolution of bite performance in *Atta*?

*Atta* ants are known for their ‘catholicity of taste’ [75, 76]; a single colony may forage on more than 100 plant species at once, spanning a large range of chemical and mechanical properties [40, 77]. Foraging is not completely indiscriminate, however: for example, foragers seem to prefer young (tender) over old (tough) leaves from the same plant [40, 42, 77, 78]. Notably, this preference disappears when pre-cut leaf fragments are offered instead [41, 42], suggesting that foraging decisions are not solely driven by plant chemistry, but also by mechanical considerations. Because larger ants typically cut tougher and denser leaves than smaller ants [5, 8, 16, 17, 40, 41], we surmise that these mechanical considerations are further confounded by size. From these simple observations emerge two key demands on bite forces in *Atta*: its magnitude must be large enough to cut a representative leaf, and its scaling determines the range of plant leaves that can be cut by workers of different size.

In order to contextualise the magnitude of the bite force, we estimate the forces required to cut leaves. Consider a blade-like tool which exerts a force of magnitude *F* to make a cut of length *dx* through a thin sheet of thickness *t*. The work done, ∫ *Fdx*, supplies the energy required to create the new surface arising from the cut, *Gtdx*, where *G* is the energy associated with a unit area of surface; the cutting force then follows as *F*_*c*_ *≈ Gt*. This calculation provides a lower bound on the required force, because it neglects the influence of friction, tool geometry and sheet bending [e. g. 79–81]. Onoda et al. used cutting tests to quantify a proxy for *G*, the work per unit fracture length to cut leaf lamina of known thickness, for about 1000 tropical plant species [82, 83, we note that although *A. vollenweideri* is often referred to as a grass-cutting ant, workers also cut dicotyledonous leaves, also see 84]. On the basis of the simple mechanical model and these extensive experimental results, we estimate that the forces required to cut tropical leaves vary between 7 - 828 mN. The median required cutting force of 82 mN can be generated by a worker of about 5 mg (see Fig. 4B), and we submit that this finding strongly suggests that the remarkable magnitude of bite forces in *A. vollenweideri* (see below) arises from the ecological need to cut leaves.

**Figure 4.**
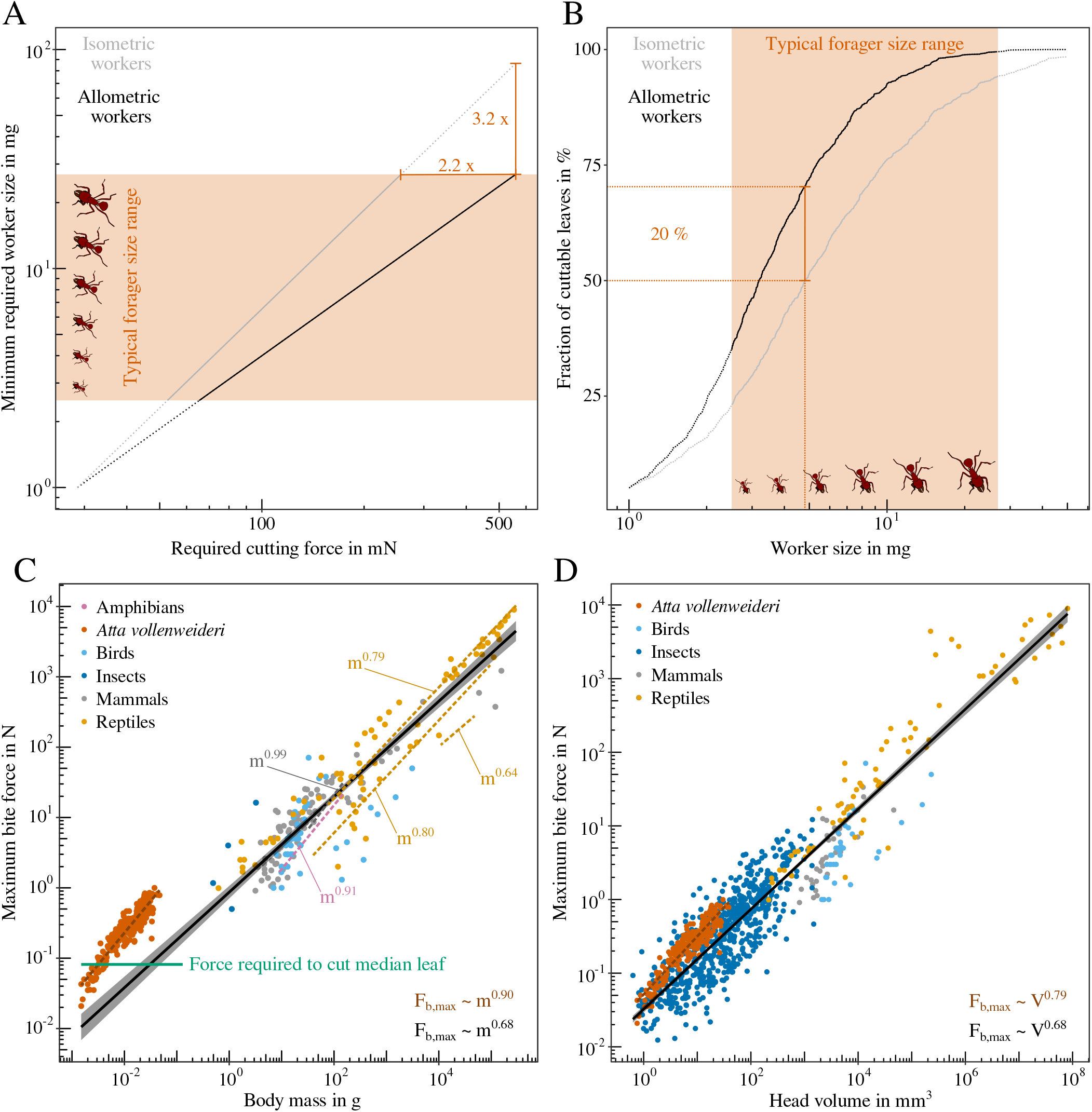
(**A**) We consider possible benefits of the positive allometry of bite force by calculating the minimum worker size which can generate a given force, both for measured positively allometric and hypothetical isometric workers. An isometric worker needs to be about three times larger to produce the same bite force as the largest allometric forager; typical *Atta vollenweideri* foragers range between 2.5 to 26.9 mg [29]. (**B**) An allometric workforce increases the total fraction of cuttable leaves, estimated from measurements on 1000 tropical plant species [83], from around 94 % to almost 100 %. A medium-sized forager of about 5 mg can cut 70 % of leaves; its isometric counterpart only 50 %. The positive allometry of bite forces thus likely increases harvesting speed, and provides flexibility in task allocation, so enhancing colony fitness. (**C**) Positive allometry of bite force has been reported in intraspecific studies for several taxa [dashed lines, see 88–92, and Table 1]. Across distantly related species and a large size range, however, bite forces scale with isometry [black line, mass^0^.^68^, OLS 95 % CI: (0.65 | 0.72), R^2^ = 0.86]; each circle represents the average maximum bite force for a single species [extracted from 45, 46, 93]. *A. vollenweideri* ants generate exceptionally large weight-specific bite forces, comparable to those of amniotes 20 times heavier. (**D**) Relative to their head volume, however, their bite forces are comparable to those of other insects [extracted from 94], and amniotes [93], suggesting that leaf-cutter ants have a large weight-specific head volume. Across species, bite forces are isometric, head volume^0.68^ [OLS 95 % CI: (0.66 | 0.70), R^2^ = 0.87]. Within *A. vollenweideri*, however, bite force ∝ head volume^0.79^ [OLS 95 % CI: (0.75 | 0.83), R^2^ = 0.86], in excess of isometric predictions, due to a disproportionate increase in volume-specific physiological cross-sectional area of the mandible closer muscle [44].

In order to contextualise the positive allometry of bite force, we next calculate the fraction of cuttable leaves as a function of worker size, both for allometric – as experimentally observed (|**F**_*b,max*_| ∝ *m*^0.90^) – and hypothetically isometric workers (|**F**_*b,max*_| ∝ *m*^0.67^, see Fig. 4). A direct comparison between two populations of workers which follow different scaling laws requires to specify the body mass at which the two scaling lines intersect. Do large workers bite relatively more strongly, or small workers relatively more weakly? This question cannot be answered *a priori*, and we thus refer to three biological arguments instead: First, the worker-size distribution in leaf-cutter ant colonies typically has a long right tail, i. e. the vast majority of foraging workers is relatively small [5, 10, 85, 86]. Second, large workers are only produced in larger numbers once colonies exceed a critical size [10, 86]. Third, larger workers are more ‘costly’ than smaller workers. On the basis of these arguments, increasing the bite force capacity disproportionally may represent a strategy to minimise the cost associated with a unit increase in maximum bite force. A disproportionate decrease of bite forces, in contrast, appears to have no obvious biological benefit. Thus, small workers may represent a reasonable generalist ‘starting point’, and we choose a body weight of 1 mg as intersection mass.

**Table 1.** Scaling of maximum bite forces for intraspecific (or intrageneric ‡) scaling studies across various taxa. The statistical models used to characterise the scaling relationships include Ordinary Least Squares (OLS) and Reduced Major Axis (RMA) regressions performed on log_10_-transformed data; in three studies, a non-linear RMA regression on untransformed data was used instead [90, 91, 95]. The relevant morphological variables are body mass (m), or one of several characteristic lengths, L: manus length (ML), snout vent length (SVL), carapace length (CL), and body length (BL). Where possible, we selected body mass as key variable to facilitate comparison with this study. A range of scaling coefficients is given when more than one species (or genus ‡), intraspecific morphotypes or sexual dimorphism were studied. Significant deviations from isometric scaling, *i* (m^0.67^, L^2^) are indicated with a *p* for positive allometry or *n* for negative allometry. We calculated a log_10_-transformed bite force quotient (BFQ) based on the reported mass range and regression coefficients [see 61, 96]. Because the BFQ is based on isometry, studies that reported allometric scaling are characterised by a range of BFQs. Where regression coefficients for body mass were missing, we calculated the range of BFQ based on the reported mass and bite force ranges (labelled with *∗*). Bite forces that were not measured at the most distal tip of the jaws, or for which the bite position was not explicitly reported, are labelled with a †; these data are biased and thus need to be compared with care (see main text).

| Taxon | Statistical model | Scaling | Allometry | $\log_{10}(\text{BFQ})$ | Source |
| --- | --- | --- | --- | --- | --- |
| Ants | OLS | $m^{0.90}$ | $p$ | 2.50 - 2.85 | this study |
| Ants | RMA | $m^{0.97}$ | $p$ | 2.44 - 2.90 | this study |
| Birds ‡ | RMA | $m^{0.76-0.88}$ | $i$ | 0.62 - 1.22 * | [97] |
| Crabs | not specified | $m^{0.82}$ | $p$ | 2.66 - 2.94 † | [98] |
| Crabs ‡ | OLS | $\text{ML}^{1.49}$ | $n$ | no mass reported | [99] |
| Crocodylians | OLS | $m^{0.79}$ | $p$ | 1.92 - 2.36 † | [88] |
| Crocodylians | RMA | $m^{0.77-0.78}$ | $p$ | 1.98 - 2.40 † | [100] |
| Frogs | RMA | $m^{0.91}$ | $p$ | 1.59 - 1.88 | [92] |
| Eels | not specified | $\text{BL}^{2.79-2.94}$ | $p$ | no mass reported † | [101] |
| Lizards ‡ | RMA | $\text{SVL}^{3.83-4.60}$ | $p$ | 1.27 - 2.34 * † | [102] |
| Lizards | RMA | $\text{SVL}^{2.65-3.37}$ | $p$ | 2.10 - 2.67 * † | [103] |
| Lizards | OLS | $m^{0.57-0.73}$ | $n, p$ | 2.02 - 2.36 * | [104] |
| Piranhas | OLS | $\text{BL}^{2.30}$ | $p$ | 2.37 - 2.47 * | [105] |
| Ratfish | OLS | $\text{BL}^{2.34}$ | $i$ | 2.07 * | [106] |
| Rodents | RMA | $m^{0.89-0.99}$ | $p$ | 1.83 - 2.04 | [89] |
| Rodents | RMA | $m^{1.11}$ | $p$ | 1.91 - 2.06 * | [107] |
| Rodents | non-linear RMA | $m^{0.93-1.01}$ | $p$ | 1.78 - 2.07 | [95] |
| Rodents | OLS | $m^{0.83}$ | $p$ | 1.78 - 2.06 * | [108] |
| Snakes | OLS | $m^{0.28}$ | $n$ | 0.71 - 1.14 † | [109] |
| Turtles | RMA | $\text{CL}^{1.73-2.23}$ | $n, i$ | 1.52 - 2.53 * † | [103] |
| Turtles | RMA | $m^{0.98}$ | $p$ | 1.90 - 2.48 † | [110] |
| Turtles | non-linear RMA | $m^{0.80}$ | $p$ | 1.40 - 1.85 | [90] |
| Turtles | non-linear RMA | $m^{0.64}$ | $i$ | 1.33 | [91] |

Viewed in this light, the positive allometry of bite force has substantial benefits for the colony for at least two reasons. First, the maximum bite force of an allometric worker with a body mass of 27 mg, at the upper end of forager sizes typically reported in *Atta* [12, 16, 17, 29, 87], exceeds that of an equally-sized isometric worker by a factor of two (see Fig. 4A). Provided that the data reported by Onoda et al. [83] are representative, this difference increases the fraction of cuttable leaves from 94 % to almost 100 % (see Fig. 4B). This difference may seem small, but this perception is erroneous: A large isometric forager would need to be three times heavier to cut the same fraction of leaves. Second, the smallest worker that can cut the median leaf almost halves in mass between isometric and allometric workers (5 to 3 mg, see Fig. 4B). As the majority of workers are typically small [86], the positive allometry thus significantly increases the fraction of the colony that can partake in leaf-cutting: an allometric forager of 5 mg can cut up to 70 % of leaves, a whopping 20 % more than its isometric counterpart. This substantial increase in the size range which can forage on a large number of plant species facilitates flexible task assignment, likely increases harvesting speed and overall influx of nutrients to the nest, and may thus be reasonably expected to enhance colony fitness.

In addition to the positive allometry of maximum bite force, our results revealed a significant decrease in the opening angle at which fibres take their optimum length, *θ*_*opt*_. We note that the last correction step, which led to this conclusion, is complex. In contrast to the first three corrections steps which are based on geometrical relations and thus have high predictive accuracy, it involves assumptions on muscle physiology, and neglects passive force-length effects due to muscle or connective tissue elasticity [47]. In support of the validity of our opening angle correction and the resulting conclusion, three independent lines of evidence may be offered. First, the fully-corrected regression analysis explains a substantially larger share of the variation of bite force with mass (86 % vs 74 % for |**F**_*b,θ*_ |, see Fig. 3B). Second, the regression residuals are independent of opening angle [ANOVA: F_1,246_ = 2.43, p = 0.12, see Fig. 3C], which does not hold without opening angle-correction [ANOVA: F_1,246_ = 148, p < 0.001]. Third, the relationship between *θ*_*opt*_ and body mass remains significant even after removing the smallest and largest size classes, which contain fewer data points (see methods), and are thus associated with higher uncertainty [OLS: slope = -0.08, 95 % CI: (−0.15 | -0.01), p < 0.05, R^2^ = 0.44]. Our result therefore appears statistically robust, and we next offer a cautious functional interpretation. Ant workers large or small may often forage on the same plant leaf. Although their size differs, the characteristic dimension of the leaf lamina – its thickness – is the same. Because cuts are typically initiated at the leaf-edge by drawing mandibles together like scissors, ants need to produce sufficiently large bite force at a gape width comparable to the leaf lamina thickness. For an ant of 3 mg, the strongest bites occur at a mandible gape of 0.1 mm, approximately half the median leaf thickness [83]. At 0.2 mm gape, the opening angle is only a few degrees larger and the force is almost identical (≈ 99 %). If, however, this small ant had the same *θ*_*max*_ as the largest colony workers, the force at 0.2 mm would be 10 % less. We thus suggest that shifts in the optimal length such that the maximum bite force occurs at larger opening angles in small workers may represent an adaptive strategy to counter size-specific disadvantages when workers of different size forage on leaves with a similar thickness.

### Positive allometry of bite force is common within, but rare across species

We discussed the mechanistic origin and ecological significance of the positive allometry of bite forces in *A. vollenweideri*. As a last step, we place our findings in a broader comparative and evolutionary context.

To this end, we compare both the scaling and magnitude of bite forces with two extensive datasets including close to 900 species, covering eight orders of magnitude in body mass and head volume [93, 94]. Regression analysis on log_10_-transformed bite force data for 203 amniote and four insect species against body mass [45, 46, 93, 111, 112], and for 139 amniote and 653 insect species against head volume [93, 94] suggests isometry of bite forces [mass^0.68^, OLS 95 % CI: (0.64 | 0.72), R^2^ = 0.86; head volume^0.68^, OLS 95 % CI: (0.66 | 0.70), R^2^ = 0.87, see Fig. 4]. The intercepts of the regression, which reflect mass- and head-volume specific bite forces (with mass in g, volume in mm^3^, and force in N), in turn are -0.06 [mass across species: OLS 95 % CI: (−0.15 | 0.03)], 1.17 [mass within *A. vollenweideri*: OLS 95 % CI: (1.07 | 1.26), see Fig. 4C], -1.49 [volume across species: OLS 95 % CI: (−1.53 | -1.44)], and -1.29 [volume within *A. vollenweideri*: OLS 95 % CI: (−1.33 | -1.25)]. Leaf-cutter ants thus appear exceptional in at least two aspects: Their bite forces grow with strong positive allometry, and their weight-specific bite forces are in substantial excess of that for the average amniote and insect, respectively [see also 47]; the largest ants produce maximum bite forces comparable to that of amniote species at least 20 times heavier (see Fig. 4C), and we have argued above that such a remarkable magnitude of bite forces probably arises from the ecological need to cut tough plant matter. Notably, both the magnitude and the scaling of the bite force of leaf-cutter ants are less remarkable relative to head volume [V^0.79^, OLS 95 % CI: (0.75 | 0.83), R^2^ = 0.86, see Fig. 4D]. This discrepancy reflects the fact that a large share of the positive allometry of bite forces in leaf-cutter ants is achieved by a positive allometry of head volume [see 44], and that leaf-cutter ants, and perhaps insects in general, have relatively larger heads than vertebrates; indeed, the weight-specific head volume for *A. vollenweideri* is 671±74 mm^3^/g compared to 209±134 mm^3^/g for amniotes [see Fig. 4 and 93].

In the above argument, we have conflated evolutionary allometry involving different species, with static allometry involving individuals of the same species at identical ontogenetic stage [113]. In order to analyse if the static positive allometry of bite forces in leaf-cutter ants is indeed exceptional, and to facilitate a more appropriate comparison of the magnitude of bite forces, we collated results from 22 intraspecific and intra-generic scaling studies on bite force across twelve taxonomic groups (including this study, see Table 1. We note that the majority of intraspecific scaling studies report ontogenetic instead of static scaling coefficients). For each study, we then calculated a log_10_-transformed ‘bite force quotient’, log_10_(BFQ) = log_10_(bite force*/*body mass^2*/*3^) in N/kg^2*/*3^ [61, 96] as a measure for weight-specific bite performance. Leaf-cutter ants have the highest log_10_(BFQ) apart from coconut crabs, for which bite force were measured at a favourable mechanical advantage [98, in contrast, we report bite forces for the maximum outlever and thus small mechanical advantage].

We conclude that leaf-cutter ants are highly specialised to produce large bite forces: (i) they have large heads relative to their body mass compared to vertebrates; (ii) the volume occupation of mandible closer muscle in these heads and (iii) the geometry of the bite apparatus are close to putative theoretical optima [44, 47]; and (iv), the estimated maximum stress of the mandible closer muscle is among the highest ever measured [47].

In sharp contrast to the magnitude of bite force, the positive static allometry appears to be less remarkable: around 70 % of available intraspecific studies reported significant positive allometry of bite forces, if often less pronounced [including one study that also reported negative allometry for other phenotypes of the same species, 104]). Remarkably, and with the notable exception of bats and finches [93, 114–117], bite force studies involving different genera typically report isometric scaling [93, 118, 119, and see meta-analysis above], or even modest negative allometry [89, 107, 120–123]. There thus appears to be a systematic difference between ontogenetic, static and evolutionary scaling of bite forces. To visualise this contrasting tendency, we superimpose a small selection of scaling relationships on the overall evolutionary slope (dashed lines in Fig. 4C). The difference between ontogenetic, static and evolutionary allometry of bite forces is reminiscent of ‘transpositional allometry’ [124–126], where group means fall onto one regression line, but within-group variation is governed by a different growth law – a result which appears to be common in biomechanical scaling studies [127–129]. Positive allometry of bite forces is likely associated with direct ecological advantages, related for example to the quantity and quality of accessible food sources [e. g. 88, 103, this study]. However, increased evolutionary rates of adaptive change in bite forces, required to drive departures from isometry, appear to be the exception rather than the norm, at least across amniotes [93].

Four factors may contribute to this seeming discrepancy: First, intraspecific and intrageneric studies on bite performance may be biased toward taxa which may be reasonably expected to face strong ecological or behavioural demands on their bite forces, so increasing the probability of departures from isometry [also see 130]. Second, strong positive allometry across many decades of mass is challenging if not impossible to achieve without fundamental ‘re-design’ [131]; as an illustrative example, a hypothetical leaf-cutter ant with a body mass of 3 g would have a closer muscle with a volume that exceeds that of its head capsule. Third, in particular ‘static’ scaling studies may often be limited by comparatively narrow size ranges [e. g. 123, 132], which increases the influence of biological variation and thus decreases the accuracy of the estimated allometric slope. Fourth, few of the available scaling studies accounted for potential size-specific biases in bite force measurements (see methods), and may thus report under- or overestimates of the allometric slope. Ultimately, the differences between ontogenetic, static and evolutionary allometry of bite forces reflect species-specific needs, coupled with complex developmental, evolutionary and ecological constraints [e.g. 66, 113, 130, 131, 133–138]. Untangling the influence of these different factors constitutes an exciting avenue for comparative and evolutionary work across disparate taxa, and a large range of body sizes.

## Conclusion and outlook

Bite forces in leaf-cutter ants scale with strong positive allometry, driven and accurately predicted by morphological and physiological adaptations of the bite apparatus. The positive allometry enables the colony to access an increased number of plant species, and is thus ecologically meaningful. Our study adds strong quantitative evidence in support of the hypothesis that size-polymorphism and the associated variation in shape broadens a colony’s access to food plants [2, 5, 86, 139, 140]. However, we acknowledge that the exact relationship between worker size, food plants and cutting behaviour is complex, and is influenced by nest distance [13], mandibular wear [38], and stridulation of the ant gaster, which can reduce force fluctuations during cutting [37, 141]. To fully integrate the mechanical results of this study with foraging behaviour, direct measurements of cutting forces with differently-sized mandibles as well as behavioural assays with materials of varying mechanical properties are needed; both are currently under way in our laboratory. We hope that such work will increase our understanding of the complex interactions between polymorphism, bite force allometry and foraging ecology in leaf-cutter ants.

## Supporting information

Bite force raw data

## Nomenclature

*A*_*phys*_: Physiological cross-sectional area of the mandible closer muscle
*α*: Misalignment angle between **F**_*b,m*_ and **F**_*b*_
*β*: Muscle force-length shape parameter
**e**_1_, **e**_2_, **e**_3_: Local head coordinate system
Γ: Correction term for differences in lever arm around the beam pivot (correction ii)
**F**_*b*_: Applied bite force
**F**_*b,m*_: Measured bite force
**F**_*b,θ*_: Maximum bite force at opening angle *θ*
**F**_*b,max*_: Maximum bite force at an equivalent mandibular opening angle, *θ*_*max*_
**F**^*∗*^: Bite force-opening angle relationship, normalised with its maximum
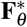: Relative bite force at opening angle *θ*
*L*_*f*_: verage muscle fibre length
**L**_*i,e f f*_: Effective mandible inlever
**L**_o,c_: Mandible outlever at bite contact point
**L**_*o,e f f,c*_: Effective mandible outlever at bite contact point
**L**_*o,d*_: Most distal (largest) mandible outlever
**L**_*o,e f f,d*_: Most distal (largest) effective mandible outlever
*L*_*opt*_: Optimal muscle fibre length at which muscle stress is maximal
*L*_*p*_: Lever arm length around the beam pivot during biting experiment
*L*_*p,cal*_: Lever arm length around the beam pivot during sensor calibration
*φ*: Average fibre pennation angle
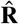: Rotational axis of the mandible joint
**S**_*l*_: Vector connecting joint centre with left head spike
**S**_*r*_: Vector connecting joint centre with right head spike
*σ*: Muscle stress
*σ*_*max*_: Maximum muscle stress
*θ*: Mandibular opening angle
*θ*_*max*_: Mandibular opening angle at which bite force is maximal
*θ*_*opt*_: Mandibular opening angle at which muscle stress is maximal

## Acknowledgments

We thank Peter Rühr and Alexander Blanke for their support on the validation of our bite force set-up, Daniela Römer for her generous support during the time of experiments, and Yusuke Onoda for sharing the leaf data. This study is part of a project that has received funding from the European Research Council (ERC) under the European Union’s Horizon 2020 research and innovation programme (Grant agreement No. 851705), a European Collaborations Grant by Imperial College (both to DL), and a Company of Biologists travel grant to FP.

