## Supplementary Materials for "Strong positive allometry of bite force in leaf-cutter ants increases the range of cuttable plant tissues"

### Effects of bite plate deflection during biting

During bite force measurements, maximum forces of around $F = 1\,\mathrm{N}$ were applied onto the bite plates. We estimate the magnitude of plate deflection caused by these forces, $\delta_{max}$, by invoking a result from simple beam theory: $\delta_{max} = FL^3/EHT^3$, where $L$ is the protruding length of the bite plate (1 mm), $H$ its height (6 mm), and $T$ its thickness ($\approx 0.15$ mm). Using a typical Young's modulus for stainless steel ($E \approx 200\,\mathrm{GPa}$), the deflection is at most $\delta_{max} = 0.0002$ mm. We hence neglected any effects of plate deflection.

### Force sensor circuit and user interface

A custom-built force set-up was developed to measure bite forces of ants and other small insects. The bite forces were measured using a capacitive force sensor, connected to a raspberry pi via $I^2C$ board (see Fig. 1A). The pi is also connected to a stepper motor via driver board, to adjust the distance between two bite plates, a camera module to record the bite experiments simultaneously to the sensor readings, and a LCD touchscreen (see main text). A user interface, written in Python (v 3.5) enables the operator to calibrate the force sensor, start and stop the force and video recordings, set the current sensor reading to zero, display live outputs of force sensor and camera, save the measurements, and to adjust the plate distance.

### Projection of the mandibular joint rotational axis

We determined the mandibular joint axis of rotation $\hat{\mathbf{R}}$ via kinematics analysis of the mandible motion [V Kang, F Püffel and D Labonte, in preparation, 1]. We assumed that the relative orientation of $\hat{\mathbf{R}}$ to the head coordinate system of the scans, defined in [2], is size-invariant. However, the relative position of $\hat{\mathbf{R}}$ with respect to the head coordinate system defined for biting ants, may differ, because it is difficult to find the same key locations for all bite force recordings. We thus defined a second coordinate system based on the coordinates of the mandible joint and the two distinct head spikes (see main text). To quantify the variation of $\hat{\mathbf{R}}$ due to difference in spike location across sizes, $\hat{\mathbf{R}}$ was projected from the scan coordinate system to the spike-based coordinate systems of all scans. The angular difference between the projections of $\hat{\mathbf{R}}$ and their mean was small, $4 \pm 1°$ with no significant size effect (Pearson's correlation coefficient: $r_{11} = 0.46$, P = 0.12). We hence selected the mean orientation of $\hat{\mathbf{R}}$ with respect to the head spikes for all sizes for the bite video analysis (see main text).

### Filament length

The length of the filaments $L_{fil}$ connecting muscle fibres and apodeme determines the relationship between muscle fibre length and mandibular opening angle (see below). We estimated $L_{fil}$ as the difference between the internal radius and apodeme radius, $L_{fil} = r_i - r_{apo}$. The internal radius represents the effective radius of attachment; the apodeme radius was defined as the equivalent radius of the apodeme cross-sectional area, assuming a circular shape, [see 2, for an exact definition].

### Variation of morphological force determinants with opening angle

In order to correct the bite force measurements in this study for size-dependent differences in opening angle, we predicted the variation of bite force across the entire worker size range, based on a previously derived biomechanical model [3]. To this end, we extracted reference measurements from tomographic scans, and used the following equations, as derived in Püffel et al. [3], in combination with Eq. 3 in the main text:

The effective inlever length, $|\mathbf{L}_{i,eff}(\theta)|$, varies with opening angle, $\theta$, as:

$$|\mathbf{L}_{i,eff}(\theta)| = \sin(\theta_0 - \theta + \gamma_0)|\hat{\mathbf{R}} \times \mathbf{L}_i||\hat{\mathbf{R}} \times \hat{\mathbf{A}}| \tag{1}$$

where $\gamma_0$ is the apodeme angle at a reference angle, $\theta_0$; $\mathbf{L}_i$ is the mandible inlever; $\hat{\mathbf{R}}$ is the joint axis of rotation; and $\hat{\mathbf{A}}$ is the apodeme main axis. The apodeme displaces as:

$$\Delta(\theta) = [\cos(\gamma_0) - \cos(\theta_0 - \theta + \gamma_0)]\frac{|\hat{\mathbf{R}} \times \mathbf{L}_i|}{|\hat{\mathbf{R}} \times \hat{\mathbf{A}}|} \tag{2}$$

The fibre pennation angle varies with $\theta$ as:

$$\phi(\theta) = \arctan\left(\frac{\sin\phi_0}{\cos\phi_0 - \Delta(\theta)/L_{t,0}}\right) \tag{3}$$

where $L_{t,0}$ is the average total length of the muscle fibres (sum of fibre length and filament length) at $\theta_0$. Filament-attached fibres vary in length as:

$$L_f(\theta) = \sqrt{\begin{array}{l}[\cos\phi_0(L_{f,0} + L_{fil}) - \Delta(\theta)]^2 \\ +[\sin\phi_0(L_{f,0} + L_{fil})]^2\end{array}} - L_{fil} \tag{4}$$

where $L_{f,0}$ is the average length of filament-attached fibres. Fibre length changes, in turn, affect muscle stress and thus bite force (see Fig. 1G and main text). For more details on parameter definitions and the geometrical basis of these equations, see Püffel et al. [3].

Table 1 | Results of reduced major axis regressions describing the relationship of bite force with body mass and head volume (labelled with ∗), respectively, on $\log_{10}$-transformed data. 'Fully-corrected' implies that all four correction steps were done (see main text), 'angle-uncorrected' means that corrections (i-iii) were performed, and 'raw' refers to uncorrected data. 95 % confidence intervals are provided in parentheses.

| Description | Units | Elevation | Slope | $R^2$ |
|---|---|---|---|---|
| Morphological prediction | mN, mg | 1.41 (1.31, 1.51) | 0.93 (0.84, 0.1.03) | 0.98 |
| Fully-corrected measurement | mN, mg | 1.38 (1.34, 1.43) | 0.97 (0.93, 1.02) | 0.86 |
| Opening angle-uncorrected measurement | mN, mg | 1.12 (1.05, 1.19) | 0.99 (0.93, 1.06) | 0.74 |
| Raw measurement | mN, mg | 1.33 (1.27, 1.39) | 0.91 (0.86, 0.97) | 0.75 |
| Fully-corrected measurement ∗ | N, $mm^3$ | -1.34 (-1.38, -1.30) | 0.85 (0.81, 0.89) | 0.86 |
| Meta-analysis | N, g | -0.17 (-0.25, -0.08) | 0.73 (0.70, 0.77) | 0.86 |
| Meta-analysis ∗ | N, $mm^3$ | -1.59 (-1.63, -1.54) | 0.91 (0.86, 0.97) | 0.87 |

Table 2 | Results of reduced major axis regressions describing the relationship of mandibular opening angle in degrees, (i) at which muscle fibre length is optimum, $\theta_{opt}$, and (ii) at which bite forces are maximum, $\theta_{max}$, with body mass in mg on $\log_{10}$-transformed data. 95 % confidence intervals are provided in parentheses.

| Quantity | Elevation | Slope | $R^2$ |
|---|---|---|---|
| $\theta_{opt}$ | 1.89 (1.83, 1.95) | -0.13 (-0.19, -0.08) | 0.56 |
| $\theta_{max}$ | 1.83 (1.79, 1.87) | -0.08 (-0.12, -0.05) | 0.59 |

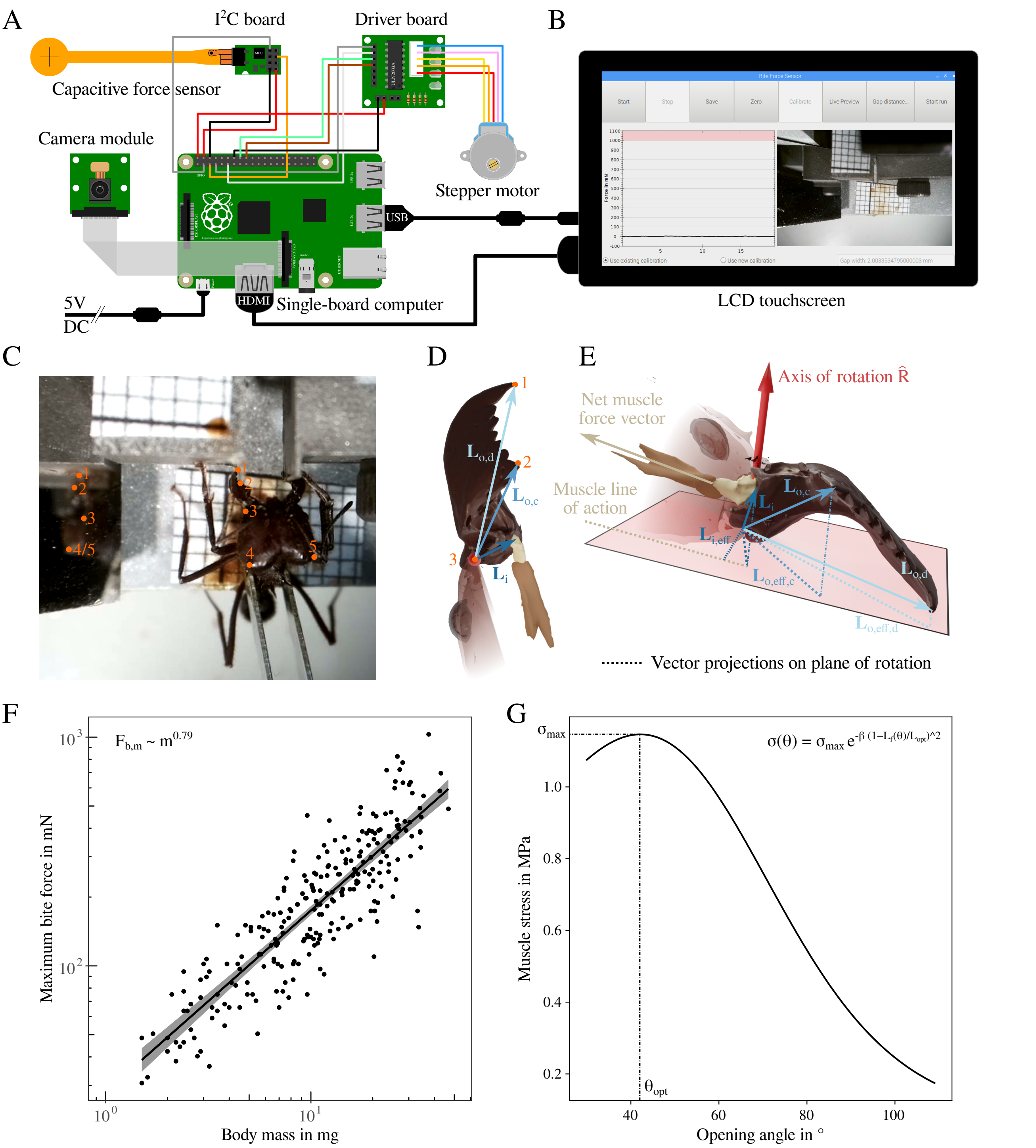


Figure 1 | (**A**) We developed a force set-up to accurately measure bite forces of small insect. Bite forces were measured by compressing a capacitive force sensor, sampled at 33 Hz (for more details, see main text). The sensor is connected to a raspberry pi via an I$^2$C board. The pi, powered at 5 V DC, is also connected to a stepper motor via driver board, a camera module, and a LCD touchscreen. (**B**) A user interface, written in Python (v 3.5), enables the experimenter to calibrate the force sensor, start and stop the force and video recordings, set current force readings to zero, show live readings of forces and camera output, save measurements, and move the stepper motor. (**C**) This force setup was used to measure bite forces of differently-sized *Atta vollenweideri* leaf-cutter ants. In order to correct the measured bite forces for size-dependent confounding effects, the coordinates of five landmarks were extracted from each video recording, as measured from top and side view: (1) most distal tooth tip, (2) bite contact point, (3) mandible joint centre, (4) left head spike, (5) right head spike. Landmarks (3-5) were used to define a local coordinate system for each ant head in order to project the joint axis of rotation onto it (for more details, see main text). (**D**) & (**E**) Landmarks (1-3) spanned the mandible outlevers, $\mathbf{L}_{o,c}$ and $\mathbf{L}_{o,d}$. We then obtained the effective outlevers, $\mathbf{L}_{o,eff,c}$ and $\mathbf{L}_{o,eff,d}$, respectively, via projection onto the plane of rotation. The mandible inlever, $\mathbf{L}_i$, measured from tomographic scans, was used to extract the effective inlever, $\mathbf{L}_{i,eff}$, defined as the shortest vector which connects the line of action of the net muscle force vector with the axis of rotation. (**F**) Without any corrections, the maximum measured bite force, $\mathbf{F}_{b,m}$, is proportional to $m^{0.79}$; this scaling coefficient is substantially lower than that obtained for fully-corrected data (0.90, see main text). (**G**) In order to correct the force data for size-dependent differences in mandibular opening angle, we used physiological parameters of the mandible closer muscle extracted for closely-related *A. cephalotes* ants [3], and fitted the opening angle at which muscle stress peaks, $\theta_{opt}$, for each of the 13 different size classes (see Eq. 3 in the main text). Maximum muscle stress, $\sigma_{max}$ and shape parameter $\beta$ were retained across species and size; $\theta_{opt}$ decreased significantly with body mass. In accordance with the results of this study, muscle stress of *A. cephalotes* majors peaks at comparatively small opening angles, $\theta_{opt} \approx 43°$ [3].
